# Selection of active defensive behaviors relies on extended amygdala dopamine D2 receptors

**DOI:** 10.1101/2021.02.24.432692

**Authors:** Laia Castell, Valentine Le Gall, Laura Cutando, Emma Puighermanal, Daniel Jercog, Pauline Tarot, Adrien Tassou, Anne-Gabrielle Harrus, Marcelo Rubinstein, Régis Nouvian, Cyril Rivat, Cyril Herry, Emmanuel Valjent

**Affiliations:** IGF, Univ. Montpellier, CNRS, Inserm, F-34094 Montpellier, France; Université de Bordeaux, Neurocentre Magendie, U1215, F-33077 Bordeaux, France; INM, Univ. Montpellier, Inserm, F-34000 Montpellier, France.; Instituto de Investigaciones en Ingeniería Genética y Biología Molecular, CONICET; FCEN, Universidad de Buenos Aires, Buenos Aires, Argentina; and Department of Molecular and Integrative Physiology, University of Michigan Medical School, Ann Arbor, Michigan, USA

**Keywords:** dopamine, central amygdala, bed nucleus stria terminalis, active avoidance, innate and learned threat, D2R

## Abstract

The ability to efficiently switch from one defensive strategy to another maximizes an animal’s chance of survival. Here, we demonstrate that the selection of active defensive behaviors requires the coordinated activation of dopamine D2 receptor (D2R) signaling within the central extended amygdala (EA) comprising the nucleus accumbens, the oval bed nucleus stria terminals and the central amygdala. We find that discriminative learning between predictive and non-predictive threat auditory stimuli is unaltered in mice carrying a temporally-controlled deletion of D2R within output neurons of the EA. In contrast, intact EA D2R signaling is required for active avoidance learning and innate flight responses triggered by a visual threat stimulus (looming). Consequently, conditional D2R knockout mice biased defensive responses toward passive defensive strategies. Altogether, these findings identify EA D2R signaling as an important mechanism by which DA regulates the switch from passive to active defensive behaviors, regardless whether of learned or innate threat.

## Introduction

Defensive behaviors that include passive (freezing, fainting) or active (flight and fight) coping strategies are evolutionary conserved reactions elicited in response to stressful and threatening situations^1, 2^. The selection of the appropriate defensive responses depends on several factors including the nature of the threat, the context and the individual’s internal state^3^. It also relies on an animal’s ability to efficiently switch from one defensive strategy to another, thus maximizing its chance of survival^4^. Therefore, while attacks eliciting passive defensive reactions in predators are often followed by flight responses, fainting strategies generally take place when freezing, flight or fight are no longer an option^5^.

Distinct brain circuits are recruited to engage passive (i.e. freezing) or active (i.e. avoidance) coping strategies following the presentation of discrete auditory cues predicting a threat^6, 7^. Interestingly, these neural circuits converge onto output nuclei of the central extended amygdala (EA), a large forebrain unit comprising the central amygdala (CeA), the bed nucleus of the stria terminalis (BNST) and the nucleus accumbens (Acb) involved in different components defensive strategies^8–12^. For instance, activation of basolateral amygdala (BLA) to Acb pathway favors active responses such as avoidance^6^. In contrast, passive defensive behaviors can be elicited by the activation of BLA principal cells projecting to CeA^6^. Moreover, a CeA neural circuit controlling behavioral responses’ switch from passive to active strategies has been identified^5,13^ suggesting that distinct interconnected EA neural circuits may influence the selection of appropriate defensive behaviors.

Output nuclei of the central EA are tightly modulated by dopamine (DA) inputs arising from the ventral tegmental area (VTA)^14^. Increasing evidence suggests that distributed EA- projecting DA neurons encoding reward prediction error and conveying salient signals coordinately ensure the selection of the appropriate defensive strategy^15–17^. Although distinct dopamine receptors subtypes may modulate different aspects of defensive behaviors, D2 receptor (D2R) signaling has been shown to participate in the control of both passive and active coping strategies^18–20^. However, whether the switch from one defensive strategy toward another relies on coordinated EA D2R signaling remains unknown.

To address this issue, we took advantage of newly generated D2R conditional knock-out mice allowing the temporally-controlled D2R deletion selectively in the central extended amygdala. In this study, the analysis of active and passive learned and innate defensive behaviors suggests that disruption of EA D2R signaling biases defensive responses toward passive coping strategies.

## Materials and Methods

### Animals

C57Bl/6J (Charles River Laboratories, France) were used for *in situ* hybridization and *Drd2-*eGFP (D2R-eGFP) for immunofluorescence analysis. The deletion of *Drd2* from Wfs1 neurons was carried out by crossing heterozygous *Wfs1-CreERT2* mice with homozygous *Drd2^loxP/loxP^* mice^21^. For all the experiments, male mice homozygous for *Drd2^loxP/loxP^* expressing CreERT2 under the promotor of *Wfs1* gene were compared with controls (Cre-negative mice). Animals were housed under standardized conditions with a 12 h light/dark cycle, ad libitum food and water, stable temperature (22 ± 2 °C) and controlled humidity (55 ± 10%). Housing and experimental procedures were approved by the French Agriculture and Forestry Ministry (A34- 172-13). Experiments were performed in accordance with the animal welfare guidelines 2010/63/EC of the European Communities Council Directive regarding the care and use of animals for experimental procedures.

### Tamoxifen injections

Tamoxifen (100 mg/kg) was purchased from Sigma-Aldrich and administered intraperitoneally at 8 weeks of age in a volume of 10 ml/kg during 3 consecutive days. Tamoxifen was dissolved in sunflower oil and ethanol (10:1) to a final concentration of 10 mg/ml.

### Immunofluorescence

Tissue preparation and immunofluorescence analyses were performed as described in^22^. Briefly, free-floating sections (30 µm) were rinsed in Tris-buffered saline (TBS; 0.25 M Tris and 0.5 M NaCl, pH 7.5), incubated 15 min in 0.2% Triton X-100 in TBS, and blocked for 1 hr in 3% bovine serum albumin (BSA) in TBS. Slices were then incubated in 0.15% Triton X-100 and 1% BSA in TBS overnight at 4°C with the primary antibodies, chicken anti-GFP (1:1000, Life Technologies, #A10262) and rabbit anti-Wfs1 (1:500, Proteintech, 11558-1-AP). The following day, slices were rinsed three times in TBS and incubated 45 min with goat Cy3-coupled anti-rabbit (1:500; Jackson ImmunoResearch Laboratories) and goat Alexa Fluor 488-coupled anti-chicken (1:500; Invitrogen) secondary antibodies. Sections were rinsed twice in TBS and twice in 0.25 M Tris-buffer before mounting in 1,4-diazabicyclo-[2.2.2]-octane (DABCO, Sigma-Aldrich). Fluorescent images of labeled cells in the Acb, CeA and BNSTov were captured using sequential laser scanning confocal microscopy (Leica SP8).

### In situ hybridization

Mice were sacrificed by cervical dislocation, brains were removed and placed on dry ice for 5 min and then stored at -80°C. Tissue was included in an embedding medium to ensure optimal cutting temperature and then sectioned on the cryostat at 14 µm with a chamber temperature of -17°C and the object at -18°C. CeA and BNST slices were collected onto Superfrost Plus slides (Fisher Scientific). Staining for *Drd2* and *Wfs1* mRNAs were preformed using single-molecule fluorescent *in situ* hybridization (smFISH). RNAscope Fluorescent Multiplex labeling kit (ACDBio catalog #320850) was used to perform the smFISH assay according to manufacturer’s recommendations. Probes used for staining are Mm-Drd2-C3 (ACDBio catalog #406501-C3), Mm-Sst-C1 (ACDBio catalog #406631-C1) and Mm-Wfs1-C2 (ACDBio catalog #500871-C2). After incubation and amplification of the fluorescent-specific signal, slides were counterstained with DAPI and mounted with ProLong Diamond Antifade mounting medium (Thermo Fisher Scientific catalog #P36961). Fluorescent images of labeled cells were captured using sequential laser scanning confocal microscopy (Leica SP8).

### Auditory brainstem response recordings

Auditory brainstem responses (ABR) were carried out under anesthesia with Rompun 2% (3 mg/kg) and Zoletil 50 (40 mg/kg) in a Faraday shielded anechoic soundproof. Rectal temperature was measured with a thermistor probe and maintained at 38.5°C ± 1 using a heater under-blanket (Homeothermic Blanket Systems, Harvard Apparatus). The acoustical stimuli consisted of 9-ms tone bursts, with a plateau and a 1-ms rise/fall time, delivered at a rate of 11/sec with alternate polarity by a JBL 2426H loudspeaker in a calibrated free field. Stimuli were generated and data acquired using Matlab (MathWorks) and LabView (National Instruments) software. The difference potential between vertex and mastoid intradermal needles was amplified (2500 times, VIP-20 amplifier), sampled (at a rate of 50 kHz), filtered (bandwidth of 0.3-3 kHz), and averaged (100 to 700 times). Data were displayed using LabView software and stored on a computer (Dell T7400). ABR thresholds were defined as the lowest sound intensity that elicits a clearly distinguishable response.

### Mechanical and thermal sensitivity

Tactile withdrawal threshold was determined by the up-down method described by Chaplan et al.^23^. Briefly, calibrated von Frey filaments (Stoeling, Wood Dale, IL, USA) were applied perpendicularly to the plantar surface of the hindpaw in logarithmically spaced increments ranging from 0.04 to 8 g. The 50% paw withdrawal threshold was determined in grams by the Dixon nonparametric test^24^. The protocol was repeated until three changes in behavior occurred. To assess heat sensitivity, a radiant heat source (plantar test Apparatus, IITC Life Science, Woodland Hills, USA) was focused onto the plantar surface of the paw. The paw withdrawal latency was recorded. Each paw was tested 3 times with 10 min-intervals between each trial. A maximal cut-off of 20 sec was used to prevent tissue damage.

### Auditory fear conditioning

The experiment was carried out in a fear conditioning apparatus (Imetronic, Pessac, France)^25^. It consists of two soundproof boxes with different contextual configurations. Box A has smooth white foam on the walls, square cage and a metal grid as the floor. Box B has dark honeycomb foam on the walls, circle cage and a metal grid as the floor. Two variants of the auditory fear conditioning were carried out. On the simplest version, the animals underwent 10 min of habituation to box A or B, cleaned before and after the session with 70% ethanol. Then, they received 5 pairings of tone (CS+, 10 sec) with an unconditioned stimulus (US: 0.6 mA scrambled foot-shock, 2 sec) coinciding with the last 2 sec of the CS+ presentation. The interval between each pairing was a random time between 35 and 60 sec. The session finished 30 sec after the end of the last CS+. 24 hrs later the animals were placed on the contrary box as for the conditioning to undergo the test and extinction sessions for three consecutive days. On these sessions, the boxes were cleaned with 1% acetic acid and, after 10 min of habituation, the CS+ was presented alone 12 times for 30 sec with an intertrial interval between 20 sec and 3 min. The session finished 30 sec after the end of the last CS+. In the second version of the auditory fear conditioning, a second CS (CS-) was added to the protocol, consisting of a tone, different from the CS+ that was not paired with the foot-shock. In order to habituate the animals to the tones, both tones were presented for 30 sec 4 times each after 5 min of exploration. On the same day, animals were placed on the same box as for the habituation and after 5 min of exploration they received 5 pairings of the CS+ (30 sec) with the foot-shock at the last sec of the CS+ (1 sec, 0.6 mA), and 5 presentations of the CS- alone (30 sec). 24 hrs later animals underwent the test and two extinction sessions on a different box during 3 consecutive days. On these sessions, animals received 12 presentations of the CS+ (30 sec) and 4 presentations of the CS-. The intertrial interval in all sessions was between 20 sec and 3 min. In the habituation and conditioning, cages were cleaned with 70% ethanol and for the test and extinction sessions with 1% acetic acid. The meaning of each tone (paring or not with the foot-shock) was counterbalanced between animals and genotypes. In both tests freezing behavior was recorded using a tight infrared frame. The threshold for considering freezing behavior was set up at 2 sec. The first 10 min of habituation were used to assess the basal freezing.

### Fear coping

The same boxes than fear conditioning were used for this test (Imetronic, Pessac, France). Mice underwent a conditioning session consisting of 2 min habituation to the box and one pairing of a tone (CS+) lasting 20 sec with 1 sec foot-shock (0.6 mA) coinciding with the last second of the tone. The session finished 30 sec after the end of the foot-shock. 24 hrs later mice underwent a test session, where 3 min habituation served to assess basal freezing and then the tone (CS+) was presented for 8 min continuously. The session finished after a random time between 20 sec and 3 min after the tone finished.

### Contextual fear conditioning

The experiment was carried out in the same fear conditioning apparatus from the auditory fear conditioning (Imetronic, Pessac, France). Animals were placed in one of the two boxes to explore the environment for 20 min. The same day, after 5 min exploring the environment, animals underwent the conditioning session during which they received 5 footshocks (1 sec, 0.6 mA; intertrial interval between 20 sec and 3 min). 24 hrs later and for 3 consecutive days, mice underwent test and extinction sessions (one per day). Animals were placed in the same box for 35 min. Freezing behavior was recorded using a tight infrared frame. The threshold for considering freezing behavior was set up to 2 sec. The first 5 min of habituation were used to assess the basal freezing.

### Active avoidance

The experiment was carried out in a soundproof shuttle box (Imetronic, Pessac, France)^25^. The apparatus is made of two equal compartments (20 x 20 cm) separated by an opened door. Both compartments have a metal grid on the floor, independent houselights and one infrared beam frame on each compartment. Mice were subjected to a habituation session consisting of 10 presentations of two different tones, 8 sec each. The two sounds were stopped as soon as the mice moved to the adjacent compartment. 24 hrs later and for 5 days, the animals were subjected to the training sessions once per day, where they received 30 presentations of the two sounds. On the CS+, the animals had 8 sec avoidance interval and 5 sec escape interval meaning that the CS+ was presented for the duration of the interval or until the mice shuttled to the other compartment. If the animal performed an avoidance (shuttling) the CS+ was terminated. However, if the animal did not avoid an unconditioned stimulus (0.7 mA foot-shock coinciding with the last 5 sec of the CS+ presentation) was delivered in the occupied compartment. Mice were exposed to a second tone (CS-) of 8 sec that was never associated with the US. During the intertrial interval the animal was free to cross between compartments. The number of avoidances was used as an index of learning.

### Visual looming

The mice were isolated in individual cages on the first handling day, and handled by the experimenter for 5 sessions prior to the behavioral testing. The behavioral apparatus was composed of a 45 x 30 cm rectangular arena, with 20 cm high walls, with a 12 cm wide roof on one side that constituted the shelter. A screen was placed over the arena, covering ⅔ of the arena surface, to display overhead stimuli. There were two cameras (30 frames per second) fixed on top of the arena to track the mouse position online to trigger the visual stimuli when the position of the mouse was at the center of the arena and to perform behavioral analyses offline. The arena, screen and cameras were inside a box covered with isolating foam. Prior to the test, mice were allowed to freely explore the arena for 8 min on the first test day, and 3 min on the following test days. Mice were tested on 2 consecutive days per week, for 2 weeks (4 sessions in total). On each session, mice were presented with 20 trials of the looming sequence. Each looming sequence was composed of 5 repetitions of the looming stimulus, consisting of a black disk expanding to a grey background from 1 cm to 25 cm in 250 ms, then staying at that size for 500 ms (750bms in total for one looming disk, 3750bms in total for the sequence, called 5L). ITI was randomized between 30 s to 180 s and the looming sequence was displayed when the position of the animal was detected inside the zone covered by the screen. The videos of the different sessions were processed with DeepLabCut. DeepLabCut was used to extract the frame by frame x and y positions of several points, notably the nose and center of the mouse. These positions were processed using Matlab to automatize data processing and the scoring of behavioral responses. These positions were then smoothed over 5 consecutive points, and used to calculate the instantaneous speed of the mouse (using the center speed) and the instantaneous speed of its nose. Both speeds were then used to score behavioral responses. Freezing responses were scored when the position of the center of the mouse and the nose speed were simultaneously inferior to a precise speed threshold (0.65 pix/frame for the center, 0.5 pix/frame, for the nose) for a minimum duration of 0.5 sec. This double speed condition was implemented to allow the exclusion of unwanted events of immobility of the center of the mouse that were not freezing responses (nose moving when the mouse was looking, grooming or sniffing around). Flight responses were scored when the speed of the center of the mouse was superior to 15 pix/frame. We also scored whether the mouse was returning or not to the shelter after the visual stimulations or not. We scored a fear response to a visual stimulus when the onset of the fear response was occurring within a 10 sec time window following the onset of the visual stimulus.

### Statistical analyses

GraphPad Prism v7.0 software was used for statistical analyses. Data are shown as the means ± SEM. For normally distributed parameters, two-way ANOVA repeated measures and student’s t test (unpaired, two-sided) were used. * p < 0.05, ** p < 0.01 and *** p < 0.001.

## Results

### Wfs1 is expressed in the EA and co-localizes with a subpopulation of D2R

Within the EA, D2R are expressed both postsynaptically and presynaptically onto DA neurons arising from the VTA and periaqueductal gray (PAG)^14, 26, 27^. To determine the role of postsynaptic D2R, we first analyzed the degree of overlap between the expression of Wolfram syndrome 1 (Wfs1) protein known to be enriched in the EA^21, 28^, and D2R using D2R-eGFP mice (**Figure 1**). The analysis of Wfs1 immunoreactivity confirmed its enrichment in the EA. Indeed, Wfs1 neurons were preferentially detected in the Acb core and shell as well as in the oval nucleus of the BNST (BNSTov) and the lateral part of the CeA (**Figure. 1a**). As previously reported^21^, ∼40% of neurons co-expressed Wfs1/GFP in Acb core (AcbC) and shell (AcbSh) (**Figure 1b**). Double immunofluorescence analysis also revealed that a majority of D2R-containing neurons of BNSTov and CEl were Wfs1-immunlabelled (∼69% and ∼75%, respectively) (**Figure 2b**). These results were confirmed by single molecule fluorescent *in situ* hybridization showing the presence of *Drd2* mRNA in BNSTov and CEl *Wfs1*^+^/*Sst*^-^ neurons (**Figure 1c**). Together, these results suggest that conditional *Drd2* knock- out mice (D2R-cKO) generated by crossing the tamoxifen-inducible *Wfs1-CreERT2* mouse line with the *Drd2^loxP/LoxP^* line^21^, represent a suitable tool to parse the role of EA D2R in threat processing.

**Figure 1:**
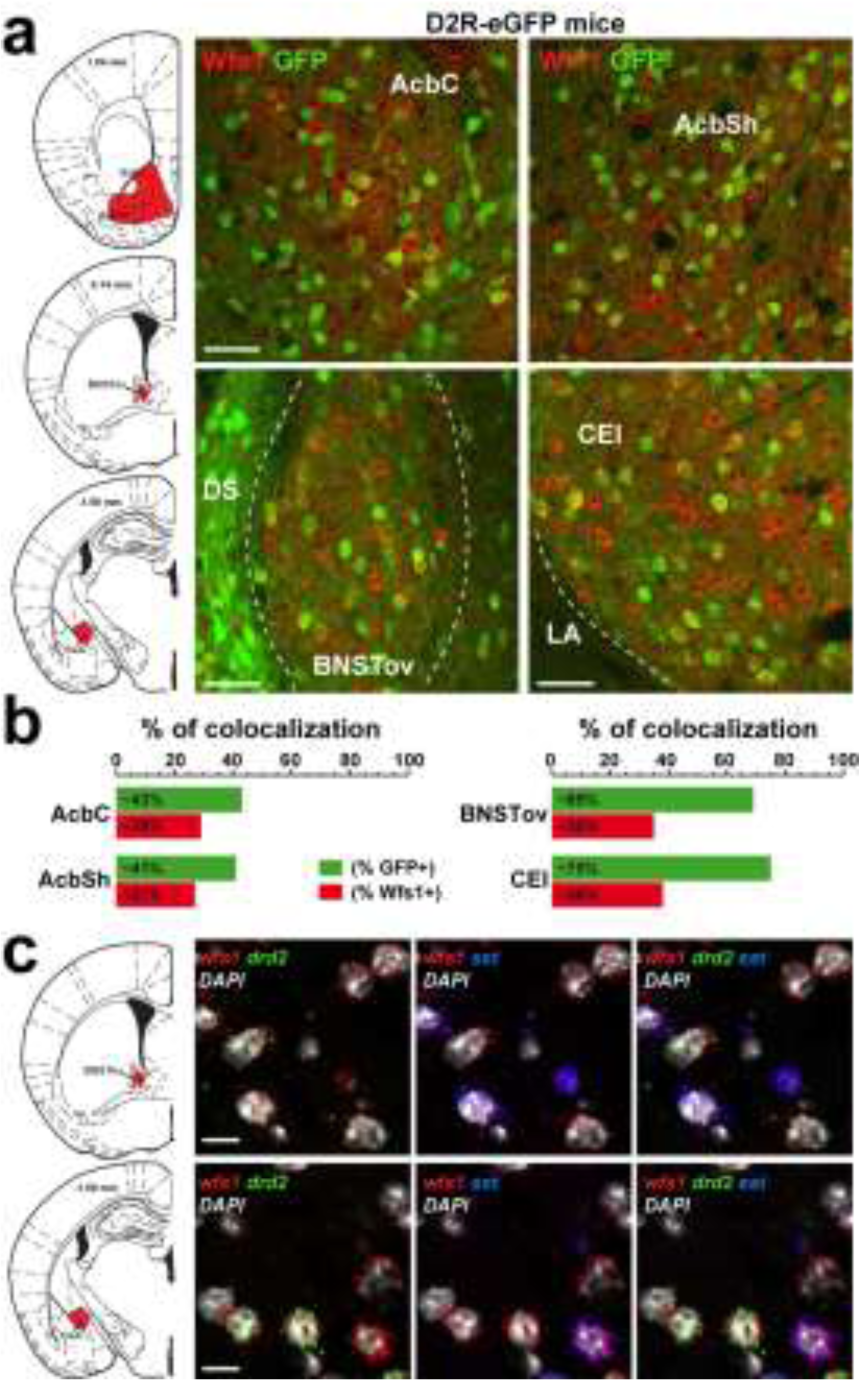
Wfs1 is expressed in the EA and co-localizes with a subpopulation of D2R neurons. (**a**) Double immunofluorescence for GFP (green) and Wfs1 (red) in D2R-eGFP mice (n = 4 mice). Scale bars: 40 µm. (**b**) Histograms showing the co-expression of the two markers as percentage of GFP-positive cells (green) and as percentage of cells expressing Wfs1 (red) in the AcbC, AcbSh, BNSTov and CEl of D2R-eGFP mice (2-3 slices per mouse, 4 mice). The total numbers of GFP- and Wfs1-positive cells counted are indicated in Table S1. (**c**) Single-molecule fluorescent *in situ* hybridization for *Wfs1* (red), *Drd2* (green) and *Sst* (blue) mRNAs in the BNSTov and CeA. Slides were counterstained with DAPI (white) (2 slices per mouse, 3 mice). Scale bars: 10 µm. AcbC: nucleus accumbens core; AcbSh: nucleus accumbens shell; DS: dorsal striatum; BNSTov: oval bed nucleus stria terminalis; CEl: lateral part of the central amygdala ; LA: lateral amygdala.

**Figure 2:**
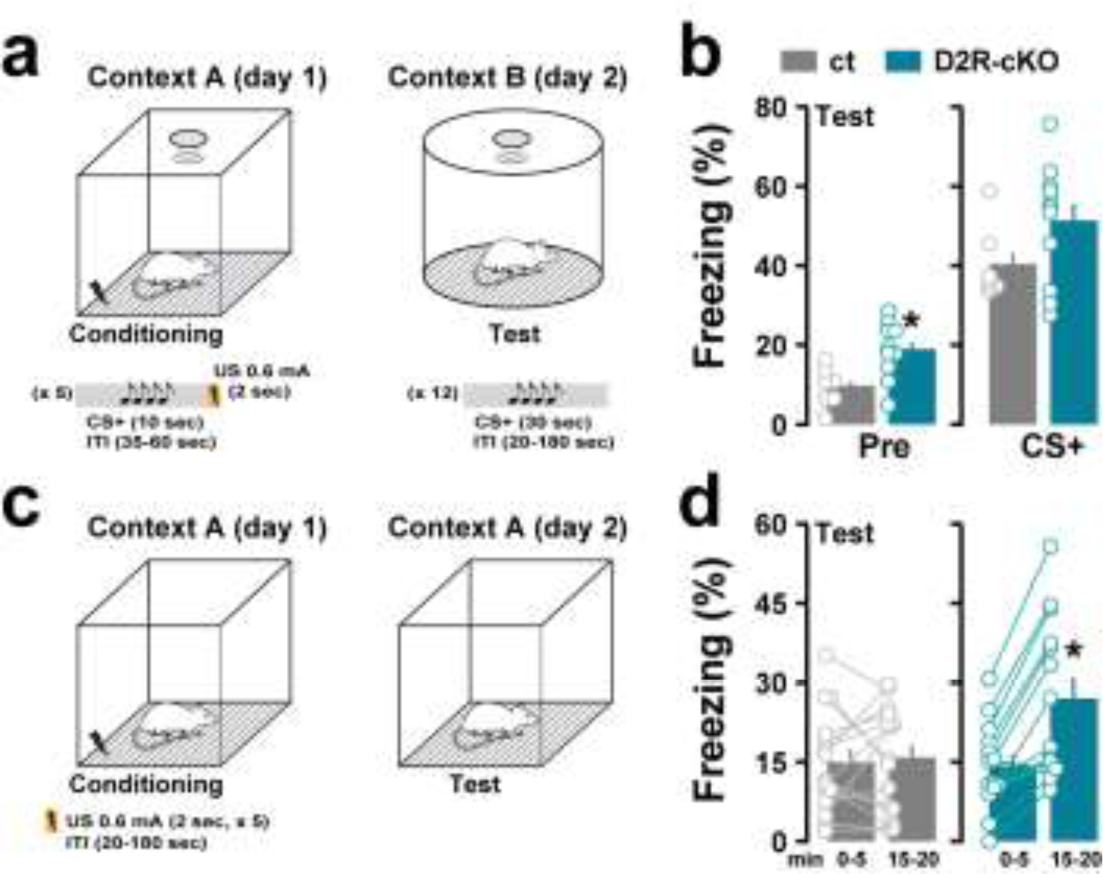
Auditory and contextual fear conditioning in D2R-cKO mice. (**a**) Schematic cartoon describing the protocol used to assess auditory fear conditioning. (**b**) Freezing responses evoked by CS+ presentation in control (grey bars) and D2R-cKO (green bars) mice. The average time spent freezing before the presentation of the CS+ (Pre) was used as a measure for contextual fear (Time: F_(1, 16)_ = 170.4, p < 0.0001; Genotype: F_(1, 16)_ = 5.399, p = 0.0336; Interaction: F_(1, 16)_ = 0.1249, p = 0.7284, two-way ANOVA followed by *post hoc* analysis Sidak’s multiple comparisons test, n = 7 ct and n = 11 D2R-cKO). (**c**) Schematic cartoon describing the protocol used to assess contextual fear conditioning. (**d**) Freezing responses evoked during the first 5 minutes and during the minutes 15-20 following the re-exposure to the context previously associated with the threat (paired *t* test: control: t14 = 0.3975, p = 0.6970; D2R-cKO: t14 = 5.763, p < 0.0001, n = 15 ct and n = 14 D2R-cKO).

### Enhanced contextual freezing responses in D2R-cKO

To determine the role of EA D2R in the control of threat processing, we first tested whether sensory systems required for learned defensive behaviors were altered in D2R-cKO mice. We focused our analysis on hearing and somatosensation which are both important when auditory cues are used as conditioned stimuli to be associated with threat (mild foot-shocks). Because Wfs1 is highly expressed in the inner ear^29^, we first measured the auditory thresholds between control and D2R-cKO mice (**Figure S1**). The audiograms were comparable between control and D2R-cKO mice (**Figure S1a**). In addition, the auditory brainstem responses, which reflect the synchronous activation of nuclei along the ascending auditory pathway, were unaltered in mice lacking D2R in WFS1 neurons (**Figure S1b, c**).

We then examined whether D2R-cKO mice displayed altered mechanical and thermal sensory thresholds. We first determined mechanical thresholds with calibrated von Frey filaments applied on both left and right hindpaws of control and D2R-cKO mice. No difference between groups was detected regarding the sensitivity to mechanical stimuli (**Figure S2**). Similar results were obtained when paw withdrawal latencies were assessed using the Hargreaves test suggesting that thermal sensitivity was also intact in D2R-cKO mice (**data not shown**). Altogether, these results indicate that processing of auditory, mechanical and thermal sensory thresholds are unaltered in D2R-cKO.

To determine the role of EA D2R in the control of threat processing, we first analyzed whether coping strategies adopted to face threatening situations were impaired in D2R-cKO mice. To do so, control and D2R-cKO mice were conditioned (context A) using a protocol during which mice learn to associate an auditory cue (CS+) with a threat (a single foot-shock delivery) (**Figure S3a**). On day 2, mice were placed in context B and exposed to the CS+ for 8 min preceded by a 2-min pre-tone period (**Figure S3b, c**). Freezing and exploration (vertical and horizontal locomotion) were measured as indexes of passive and active coping strategies, respectively (**Figure S3b, c**). During the 2-min pre-tone period, control mice displayed low freezing responses associated with high exploratory behaviors (**Figure S3b, c**). Within the two first minutes (minutes 3-5) of CS+ presentation, control mice showed strong freezing responses that gradually diminished favoring the expression of exploratory behaviors (minutes 5-8) (**Figure S3b, c**). Similar responses were observed in D2R-cKO mice suggesting that EA D2R does not play a critical role in CS-induced coping strategies.

We then examined whether EA D2R participated in the formation of auditory fear memories. Control and D2R-cKO mice underwent classical auditory-cue fear conditioning protocol (**Figure 2a**). Freezing responses were measured as the typical threat response. Control and D2R-cKO mice displaying less than 20% of freezing during the CS+ presentation were excluded from the analysis (ct n = 0 and D2R-cKO n = 1). Increased freezing responses were observed in D2R-cKO mice during the pre-tone period (**Figure 2b**). Although not significant, a tendency towards increased freezing responses was also evident in D2R-cKO compared to control mice during the CS+ presentation (**Figure 2b**). Despite this difference, both groups displayed similar freezing levels after the conditioning indicating that the ability of D2R-cKO mice to form auditory fear memories was not impaired. The enhanced freezing responses detected in D2R-cKO mice during the pre-tone period suggest however that discrete contextual cues might trigger freezing responses in D2R-cKO mice (**Figure 2b**). To test this hypothesis, we performed classical contextual fear conditioning in a different cohort of animals (**Figure 2c**). As shown in Figure 2d, control and D2R-cKO mice displayed similar low freezing responses during the first 5 minutes of re-exposure to the context in which the conditioning occurred. However, the freezing responses gradually increased over time in D2R-cKO mice while these responses were still low in control mice (**Figure 2d**). Altogether, these results indicate that EA D2R signaling modulates the expression of passive responses (i.e. freezing) to threat-conditioning contextual stimuli.

### EA D2R facilitates extinction of threat-conditioned stimulus

A previous study indicated that coordinated activation of D2R in the BNST and CeA facilitates discriminative learning between stimuli representing safety or threat^27^. We therefore evaluated whether D2R-cKO mice learn to distinguish between auditory cues associated (CS+) or not (CS-) with a threat (**Figure 3a**). During the conditioning (day 1), control and D2R-cKO mice displayed equivalent freezing responses to the CS+ and CS- (**Figure 3b**). On the test day (day 2), high levels of freezing were observed in both groups following CS+ presentation. In contrast, CS- presentation evoked low freezing responses in both control and D2R-cKO mice suggesting that discriminative learning does not require EA D2R (**Figure 3c**). We then evaluated extinction learning by presenting repeatedly the CS+ (12 times) during 2 consecutive days (day 3-4) (**Figure 3c**). While the freezing responses gradually diminished over the course of extinction in control mice, D2R-cKO mice maintained high levels of freezing which were similar to those observed immediately after conditioning (**Figure 3c-d**). Together, the results indicate that intact EA D2R signaling is necessary for extinction to occur.

**Figure 3:**
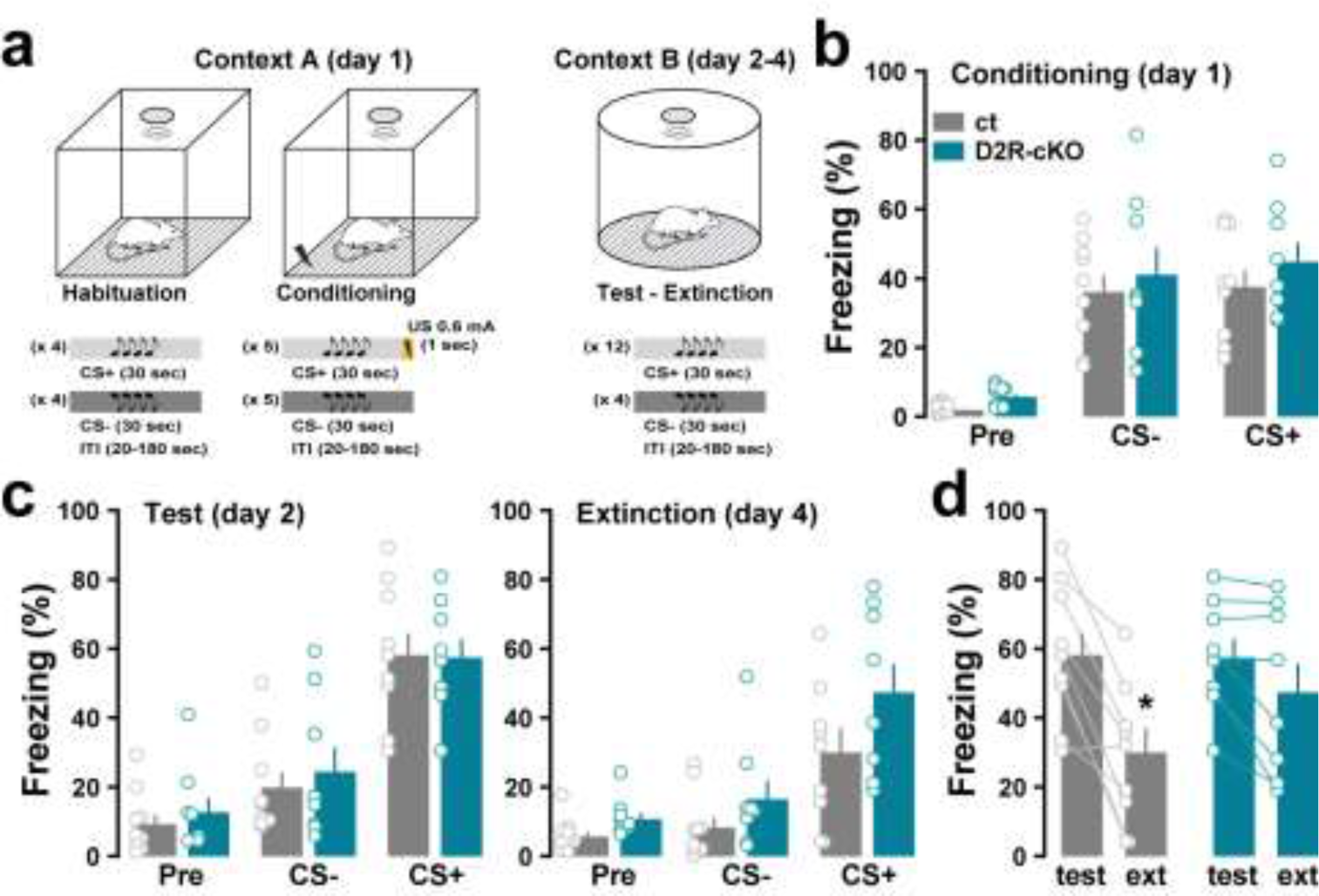
Impaired extinction learning in D2R-cKO. (**a**) Schematic cartoon describing the protocol used to assess discriminative auditory fear conditioning and extinction. (**b**) Freezing responses during the conditioning phase. Mice were exposed to CS+ and CS- cues in context A (Day: F_(2, 30)_ = 46.14, p < 0.0001; Genotype: F_(1, 15)_ = 1.140, p = 0.3025; Interaction: F_(2, 30)_ = 0.08529, p = 0.9185, two-way ANOVA). (**c**) Freezing responses evoked by the CS- and CS+ during the test (day 2) (Day: F_(2, 30)_ = 94.06, p < 0.0001; Genotype: F_(1, 15)_ = 0.1425, p = 0.7111; Interaction: F_(2, 30)_ = 0.3029, p = 0.7409, two-way ANOVA) and the extinction session (day 4) (Day: F_(2, 30)_ = 30.12, p < 0.0001; Genotype: F_(1, 15)_ = 3.671, p = 0.0746; Interaction: F_(2, 30)_ = 1.521, p = 0.2350, two-way ANOVA). (**d**) Comparison of the freezing behavior evoked by the CS+ during the test and the extinction sessions in control and D2R-cKO mice (paired *t* test: control: t16 = 3.117, p = 0.0066; D2R-cKO: t14 = 0.9845, p = 0.3416, n = 9 ct and n = 8 D2R-cKO).

### EA D2R is required for active avoidance learning

To get insights into the role of EA D2R in threat processing, we then assessed the ability of D2R-cKO mice to learn to avoid a threat by shuttling from one compartment to another during CS+ presentation (**Figure 4a**). During the first session, control and D2R-cKO mice shuttled exclusively after the delivery of shocks indicating that during this phase escape responses predominate. These data also confirm that D2R-cKO and control mice process similarly painful stimuli (**Figure 4a-b**). Upon subsequent trials, control mice gradually switched from escape to avoidance responses following CS+ presentation while D2R-cKO mice failed to do so (**Figure 4b**). Interestingly, discriminative learning was preserved in both groups as supported by the lower number of avoidance responses following CS- presentation (**Figure 4b**). Instead, the analysis of the distribution between efficient and poor learners revealed that active avoidance learning was less uniform in D2R-cKO mice compared to control mice (**Figure 4c**). Altogether, these results indicate that EA D2R are required for learned active avoidance.

**Figure 4:**
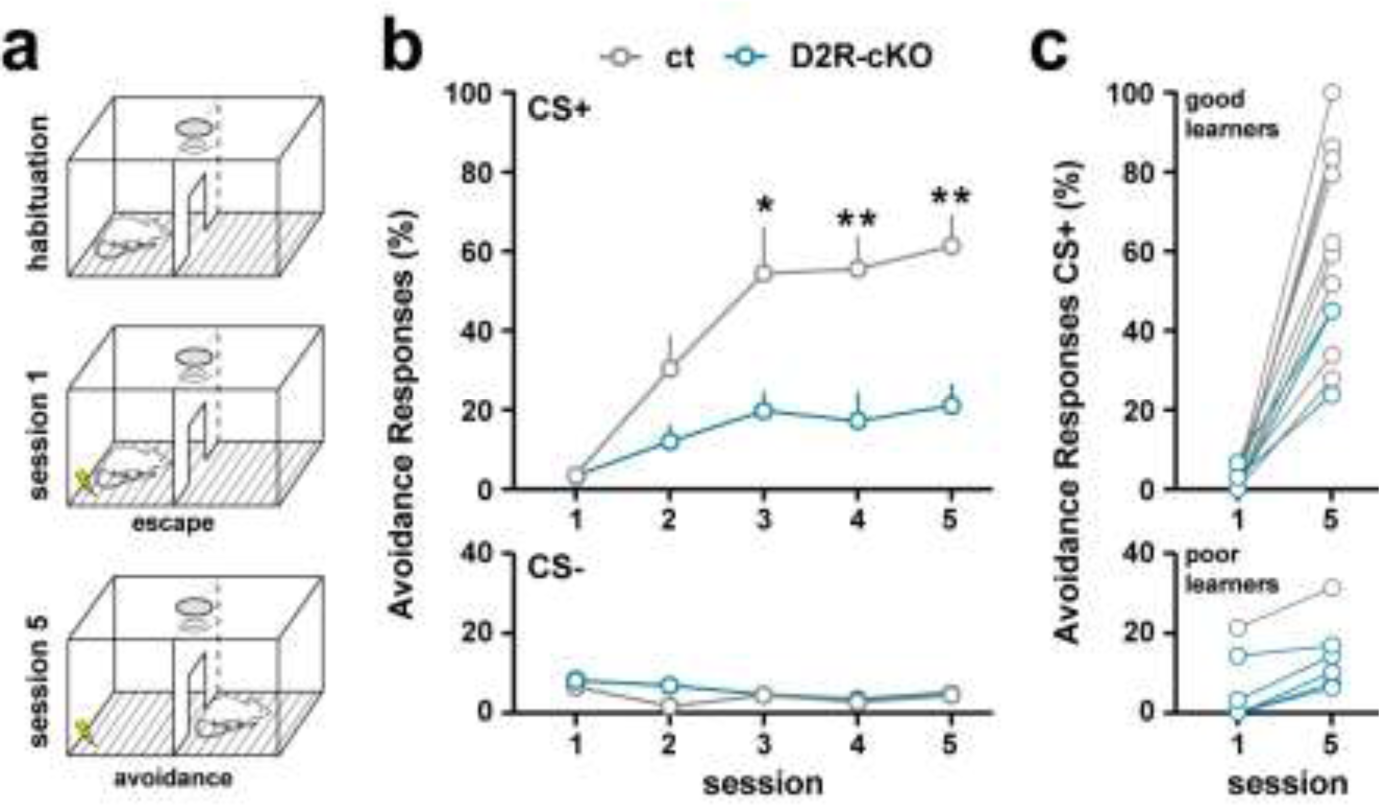
Instrumental threat learning in D2R-cKO mice. (**a**) Schematic cartoon describing the protocol used to evaluate discriminative active avoidance learning. (**b**) Avoidance responses representing the number of times the animal shuttled to the adjacent compartment during the CS+ presentation (Session: F_(5, 80)_ = 13.40, p < 0.0001; Genotype: F_(1, 16)_ = 11.92, p = 0.0033; Interaction: F_(5, 80)_ = 3.618, p = 0.0053, two-way ANOVA repeated measures) and CS- presentation (Session: F_(5, 80)_ = 3.631, p = 0.0088; Genotype: F_(1, 16)_ = 1.886, p = 0.1886; Interaction: F_(5, 80)_ = 1.220, p = 0.3075, two-way ANOVA repeated measures). (**c**) Distribution of the avoidance responses during CS+ presentation in session 1 and 5 among good and poor avoiders (n = 10 ct and n = 8 D2R-cKO).

### EA D2R biases innate defensive behaviors induced by visual threat

We next determined whether EA D2R also biases the repertoire of defensive strategies in response to innate threats. To do so, freezing and/or flight responses to looming visual stimuli thought to mimic an aerial predator were assessed in control and D2R-cKO mice (**Figure 5a**). At day 1, both control and D2R-cKO mice displayed a high probability of defensive responses (flight and/or freezing) which gradually decreased across sessions (**Figure 5b**). Moreover, both groups began their flight or freezing responses with similar latencies after stimulus onset (**Figure 5c**). However, the independent freezing and flight responses’ analysis revealed that D2R-cKO mice strongly biased their innate defensive behaviors towards freezing (**Figure 5d**). Consequently, the shelter entrance probability was strongly reduced in D2R-cKO mice compared to control animals (**Figure 5e**). Altogether, these findings indicate that EA D2R favor the selection of active defensive behaviors in response to visual innate threat.

**Figure 5:**
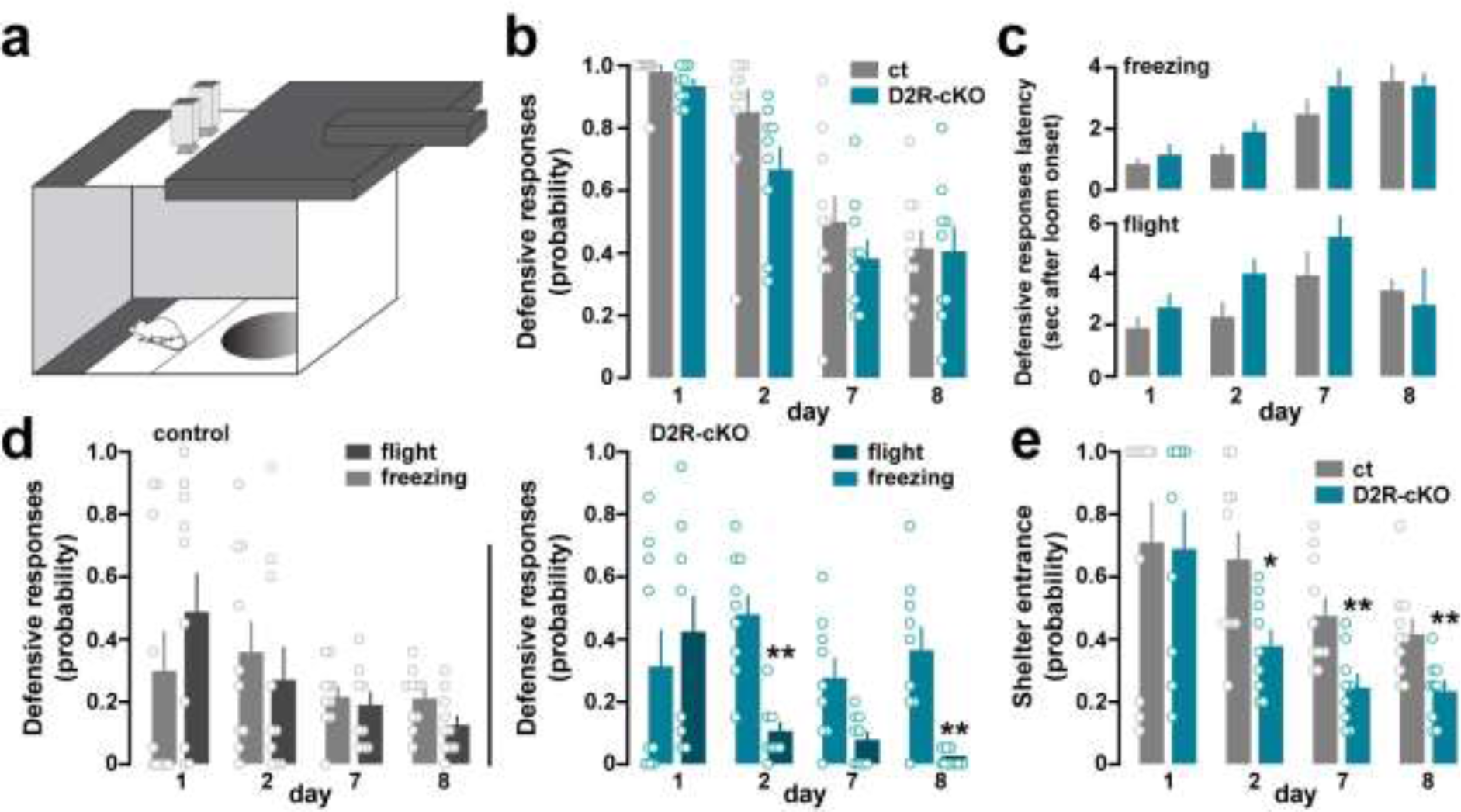
EA D2R are essential for flight responses triggered by looming visual stimulus. (**a**) Schematic cartoon describing the protocol used to evaluate the responses to the looming stimulus. (**b**) Probability of defensive responses among sessions (Session: F_(3, 51)_ = 78.97, p<0.0001; Genotype: F_(1, 17)_ =1.538 p = 0.2317; Interaction: F_(3, 51)_ = 1.582, p = 0.2051, two- way ANOVA repeated measures; n = 10 ct and n = 9 D2R-cKO). (**c**) Latency of defensive responses divided by freezing (Session: F_(3, 38)_ = 5.888, p = 0.0021; Genotype: F_(1, 17)_ = 2.757 p = 0.1152; Interaction: F_(3, 38)_ = 1.249, p = 0.3056, Mixed-effects analysis) and flight (Session: F_(3, 44)_ = 25.16, p < 0.0001; Genotype: F_(1, 17)_ = 1.641 p = 0.2173; Interaction: F_(3, 44)_ = 0.6757, p = 0.5716, Mixed-effects analysis) represented in seconds after the looming stimulus onset. (**d**) Probability of defensive responses in control (Session: F_(3, 54)_ = 4.582, p = 0.0063; Behavior: F_(1, 18)_ = 0.00078, p = 0.9780; Interaction: F_(3, 54)_ = 1.820, p = 0.1545, two-way ANOVA repeated measures) and D2R-cKO animals (Session: F_(3, 48)_ = 15.14, p = 0.0206; Genotype: F_(1, 16)_ = 11.01, p = 0.0043; Interaction: F_(3, 48)_ = 5.309, p = 0.0031, two-way ANOVA repeated measures; n = 10 ct and n = 9 D2R-cKO) divided by freezing and flight. (**e**) Probability of shelter entrance among sessions in control and D2R-cKO mice (Session: F_(3, 51)_ = 15.14, p < 0.0001; Genotype: F_(1, 17)_ = 4.771, p = 0.0432, Interaction: F_(3, 51)_ = 1.597, p = 0.2015, two-way ANOVA repeated measures; n = 10 ct and n = 9 D2R-cKO).

## Discussion

The present study investigates the role of synchronized EA D2R signaling in the control of passive and active defensive strategies induced by learned or innate threats. Our results reveal that the ability to discriminate between stimuli representing safety or threat is not impaired in mice carrying *Drd2* ablation in Wfs1 neurons from the EA. On the other hand, these mice failed to switch from passive (i.e. freezing) to active (i.e. avoidance) coping strategies following repeated presentations of cues predicting learned or innate threat and were biased toward passive coping strategies during associative learning tasks. Altogether, our findings unveil the role of EA D2R signaling in the selection of active defensive behaviors in response to both learned and innate threats.

Although the requirement of intact DA signaling in passive defensive behaviors is undeniable^30, 31^, the mechanisms by which DA regulates these behavioral responses are far from being fully understood. Our findings demonstrating that EA D2R signaling is not required for cued and contextual threat conditioning are in line with previous work performed on constitutive or striatal-specific conditional D2R deficient mice^32, 33^, but opposite to those obtained using pharmacological approaches. Thus, systemic or intra-amygdala pharmacological blockade of D2R attenuates the acquisition and the expression of conditioned freezing evoked by either context re-exposure or tone presentation previously associated with the threat^34–37^. Similar results were observed following systemic or intra-VTA activation of D2R^37–40^. These effects, which most likely result from mesoamygdala DA pathway inhibition, illustrate that D2R signaling taking place in distinct brain areas jointly but differentially participates in the control of passive defensive strategies.

Our results revealed that discrimination learning between cues representing safety or threat was intact in EA D2R-cKO mice. This suggests that generalized conditioned threat responses observed following the concomitant blockade of BNST and CeA D2R do not rely exclusively on postsynaptic EA D2R signaling, raising the intriguing possibility that sequential interplay between pre and postsynaptic D2R signaling accounts for this effect^27^. In this scenario, presynaptic filtering controlled by EA D2R signaling would contribute to select the relevant information, while postsynaptic D2R signaling would activate specific EA circuits to facilitate discriminative learning and/or select appropriate defensive behaviors. Future experiments using photoswitchable D2R ligands allowing cell type-specific and spatiotemporal control of D2R will be instrumental to test this hypothesis^41^.

Extinction of conditioned threat is classically viewed as an active inhibitory learning, characterized by a gradual decrease of the freezing responses following repeated exposure to context- or cues-predicting threats^42, 43^. Interestingly, compelling evidence indicates that the extinction process partly relies on the activation of the mesocorticolimbic DA pathway^30, 31, 44^. Indeed, DA signals conveyed by midbrain DA neurons have been shown to be necessary for and sufficient to drive extinction^45–47^. Modulation of defensive behaviors associated with extinction also requires D2R-mediated signaling. Thus, activation of D2R facilitates extinction of conditioned threat^38^ when its blockade induces extinction deficits^18, 48^ (but see^49^). Despite the prominent role of infralimbic D2R signaling in the regulation of extinction^20, 48^, our results unveiled that intact coordinated EA D2R signaling is also required thereby supporting the role of Acb D2R in extinction of conditioned threat^18^.

The extinction process is most often, if not exclusively, assessed through the prism of passive coping strategy where elevated freezing responses result from impaired extinction learning^50^. However, decision-making between passive (i.e. freezing) and active (i.e. avoidance) strategies also determines in the choice of appropriate defensive behaviors^13^. Striatal DA signaling has been implicated in the control of avoidance behaviors. Notably, Acb DA depletion disrupts operant avoidance responding^51^. Moreover, mice lacking DA displayed profound deficits in active avoidance responses, which can however be restored by normalizing DA signaling in the striatum^52^. Our findings showing that genetic deletion of D2R in EA circuits impaired conditioned avoidance responses extend previous genetic and pharmacological studies supporting a central role of D2R signaling. For example, avoidance behaviors are suppressed in mice lacking the long isoform of the D2R^53, 54^. Similar results were obtained following systemic or intra-Acb D2R blockade^55–58^. Conversely, activation of CeA D2R signaling has been shown to trigger avoidance response in poor learners^59^ further supporting the role EA D2R signaling in the selection of conditioned active avoidance. The involvement of EA D2R signaling in setting up avoidance responses to threat is however not systematic and seems to depend on the nature of the threat. Thus, the control of approach-avoidance behaviors, assessed by measuring the time spent in risky areas of the open-field (i.e. center) or the elevated plus maze (i.e. open arms), does not require the EA D2R signaling^21^, but relies on the presence of D2R in striatopallidal medium-sized spiny neurons^60^. Our work also provides evidence that intact EA D2R signaling is required for adjusting the balance between passive and active defensive behaviors in response to innate threat evoked by visual (looming) stimuli thought to mimic aerial predator approach^61^. Thus, while defensive reactions were quantitatively similar between control and EA D2R-cKO mice, the latter were biased toward freezing rather than flight, consequently failing to adapt their defensive strategies. Processing and execution of visually-evoked flight behaviors critically depend on the integrity of sequential neural pathways linking the superior colliculus (SC) to the CeA^62–66^. Interestingly, growing evidence implicates D2R signaling in SC processing and in the regulation of avoidance response to aversive stimuli^67–70^. It is however unlikely that the impaired flight response observed in EA D2R-cKO mice relies on altered SC D2R signaling since the *Wfs1* gene, which drives the Cre recombinase expression, is lacking from the SC^28^. Future studies can build on our observation to determine whether CeA D2R signaling is necessary for and sufficient to initiate flight responses evoked by threat-related visual stimuli.

In summary, our study identified EA D2R signaling as an important mechanism by which DA might regulates the switch from passive to active defensive behaviors. Additional work will establish whether the selection of defensive behaviors induced by chemical cues such as predator odors requires also the integrity of EA D2R signaling.

## Acknowledgments

The authors thank Stéphanie Trouche, Antoine Besnard, Zoé Husson and Miguel Angel Lujan for critical reading of the manuscript and their insightful comments. We thank the iExplore and MRI Platforms of the IGF for their involvement in the maintenance and breeding of the colonies. This work was supported by Inserm, Fondation pour la Recherche Médicale (DEQ20160334919), La Marató de TV3 Fundació, by the French National Research Agency ANR DOPAFEAR (EV). This work was supported by the French National Research Agency (ANR-10-EQPX-08 OPTOPATH) and the Fondation pour la Recherche Médicale (FRM- Equipes FRM 2017) (CH). Laia Castell was supported by the pre-doctoral Labex EpiGenMed («Investissements d’avenir» ANR-10-LABX-12-01). Laura Cutando was supported by the post-doctoral Labex EpiGenMed fellowship («Investissements d’avenir» ANR-10-LABX-12- 01), EP was a recipient of Marie Curie Intra-European Fellowship IEF327648 and is currently a recipient of a Beatriu de Pinós fellowship (# 2017BP00132) from the University and Research Grants Management Agency (Government of Catalonia, Spain). Anne-Gabrielle Harrus is supported by the Agence Nationale de la Recherche (ANR-15-CE16-0016-01).

## Author contribution

Laia Castell and E.V. conceived the project and designed the study. Laia Castell performed histological and behavioral experiments with the help of P.T and Laura Cutando. E.P. generated D2R-cKO mice. D.J. settled active avoidance paradigm. M.R. provided mice.

V.L.G. and C.H. performed and analyzed looming experiments. A.T. and C.R. performed and analyzed von Frey experiments. A-G.H and R.N. recorded and analyzed audiograms and auditory brainstem responses. Laia Castell and E.V. wrote the manuscript with input from all authors.

## Supplemental Figures and legends

**Supplemental Figure 1:**
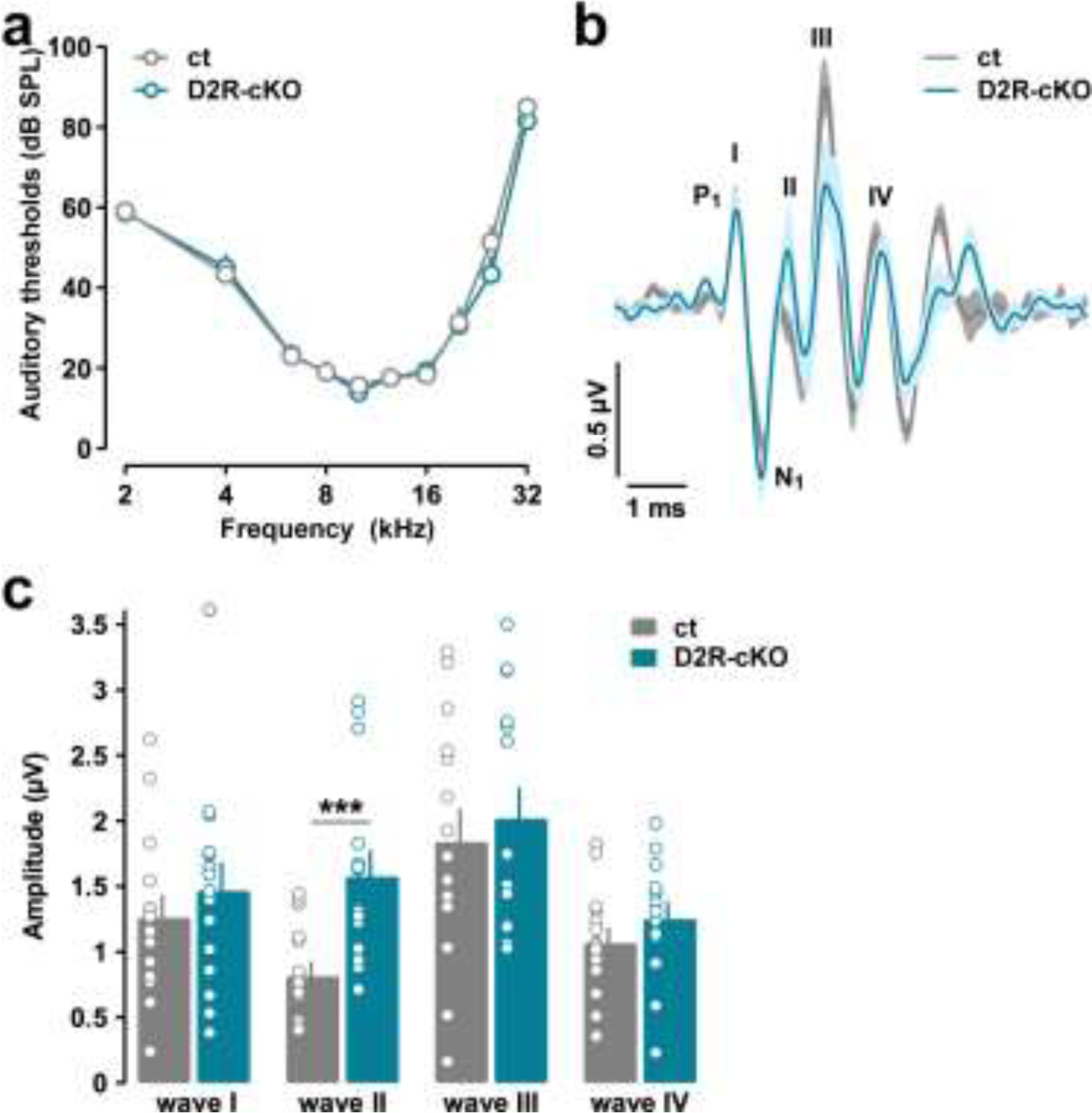
Audiograms and auditory brainstem responses in D2R-cKO mouse line. (**a**) Mean ABR audiograms from 5- to 8-month-old control and D2R-cKO mice. (Frequency: F_(9, 252)_ = 254.9, p < 0.0001; Genotype: F_(1, 28)_ = 0.2631, p = 0.6120; Interaction: F_(9, 252)_ = 0.9914, p = 0.4475, two-way ANOVA repeated measures; n = 15 ct and n = 15 D2R- cKO). (**b**) Grand average of ABR recordings (lines) and SEM (shaded areas) evoked by16- kHz tone burst at 80 dB SPL. Roman numbers indicate the Jewett ABR waves. P1 and N1 refer to the positive and negative peaks of the first wave. (**c**) Waves amplitude (positive to negative peak amplitude) average obtained from (**b**). Error bars correspond to SEM in (**a**) and (**c**). Circles represent individual measurements. Level of significance: ***p < 0.001, two- tailed Mann-Whitney Wilcoxon test.

**Supplemental Figure 2:**
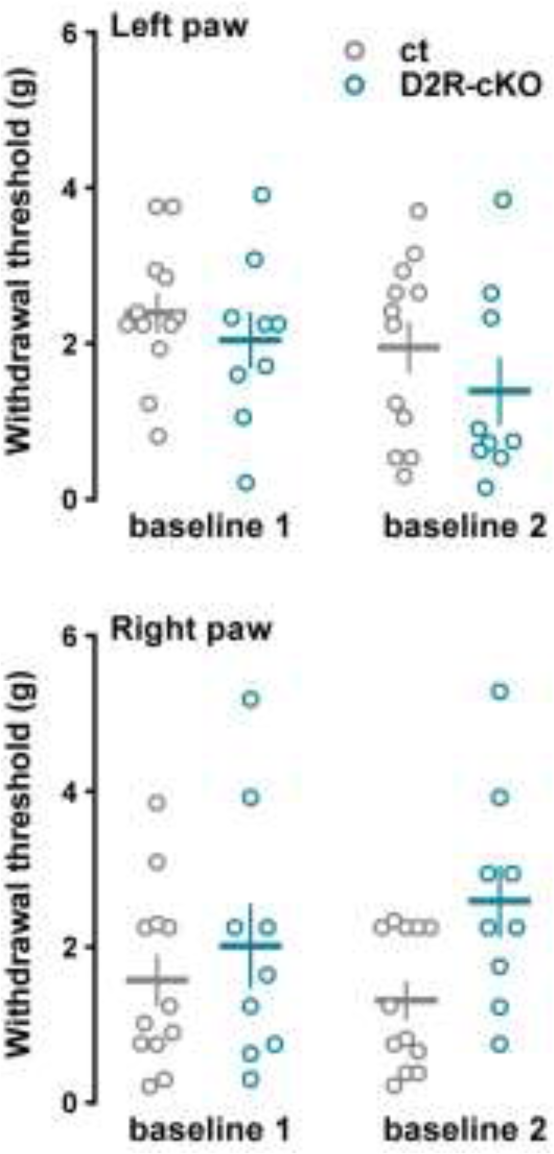
Mechanical sensory thresholds in D2R-cKO. Mechanical sensory thresholds were determined with calibrated von Frey filaments via the up and down method applied on both left (Baseline: F_(1, 19)_ = 2.643, p = 0.1205; Genotype: F_(1, 19)_ = 1.795, p = 0.1961; Interaction: F_(1, 19)_ = 0.09412, p = 0.7623, two-way ANOVA repeated measures) and right hindpaws (Baseline: F_(1, 19)_ = 0.1880, p = 0.6695; Genotype: F_(1, 19)_ = 4.192, p = 0.0547; Interaction: F_(1, 19)_ = 1.345, p = 0.2604, two-way ANOVA repeated measures; n = 12 ct and n = 9 D2R-cKO). Two independent measures were performed (baseline 1 and 2: bsl 1 and bsl 2).

**Supplemental Figure 3:**
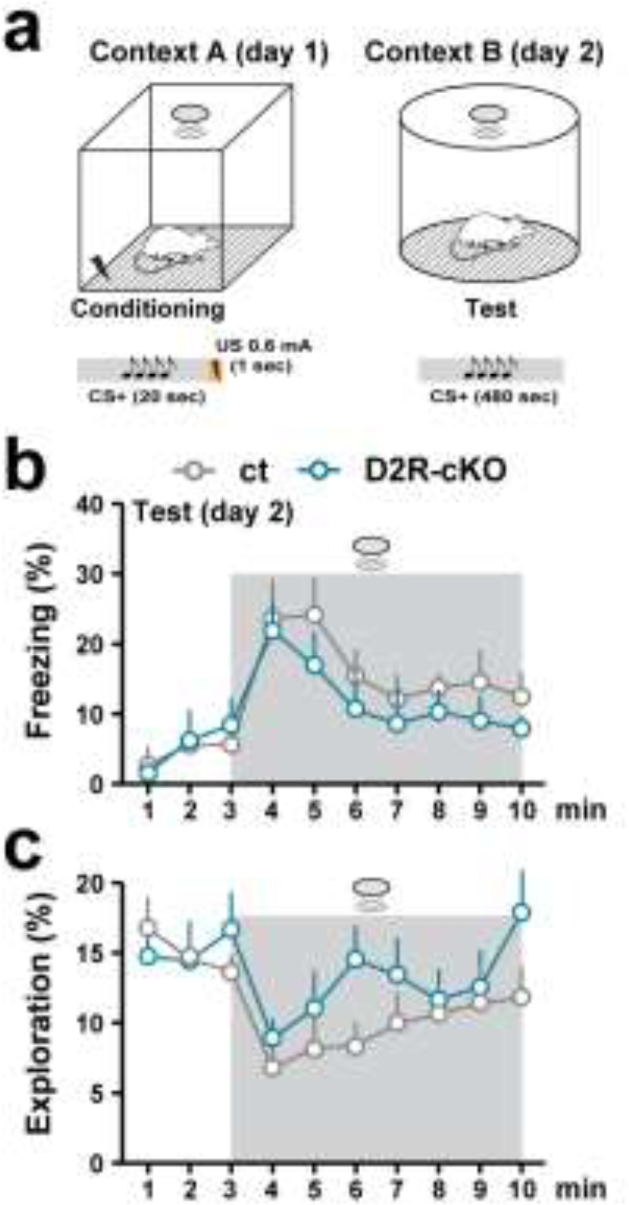
Passive and active coping strategies in D2R-cKO mice. (**a**) Schematic cartoon describing the protocol used to assess freezing and active coping responses. (**b**) Freezing response (Time: F_(9, 144)_ = 11, p < 0.0001; Genotype: F_(1, 16)_ = 0.4516, p = 0.5112; Interaction: F_(9, 144)_ = 0.6465, p = 0.7557, two-way ANOVA repeated measures, n = 9 ct and n = 9 D2R-cKO) and (**c**) exploratory behaviors (Time: F_(9, 144)_ = 6.176, p < 0.0001; Genotype: F_(1, 16)_ = 0.8530, p = 0.3694; Interaction: F_(9, 144)_ = 1.569, p = 0.1299, two-way ANOVA repeated measures; n = 9 ct and n = 9 D2R-cKO) evoked by CS+ (shaded) during the test day.

**Table S1:** Number of cells quantified in the AcbC, AcbSh, BNSTov and CeA

| Regions | AcbC | AcbSh | BNSTov | CeA |
| --- | --- | --- | --- | --- |
| GFP (total) | 512 | 496 | 299 | 457 |
| GFP (only) | 293 | 294 | 581 | 903 |
| Wfs1 (total) | 756 | 743 | 22 | 112 |
| Wfs1 (only) | 537 | 541 | 84 | 558 |
| GFP/Wfs1 | 219 | 202 | 205 | 345 |

GFP, Green Fluorescent Protein; Wfs1; Wolfram syndrome 1; AcbC; Accumbens Core; AcbSh: Accumbens Shell; BNSTov, Oval Bed Nucleus Stria Terminalis ; CeA, Central Amygdala.

